# Anthropogenic marine debris and its dynamics across peri-urban and urban mangroves on Penang Island, Malaysia

**DOI:** 10.1101/756106

**Authors:** Chee Su Yin, Yee Jean Chai, Danielle Carey, Yusri Yusup, John Barry Gallagher

**Author notes:** Corresponding author. (S.Y. Chee), (J.B. Gallagher). Declarations of interest: none.

## Abstract

Mangroves act as sinks to a variety of anthropogenic marine debris (AMD) forms. However, knowledge of their distribution and accumulation dynamics is limited. To address this shortfall, abundance, sorting, and diversity parameters of AMD were evaluated across the canopy of Penang’s urban and peri-urban mangroves. Two urban and two peri-urban mangroves were sampled at different periods over 2 months, with differences constrained by possible changes in their wind fields, and neap-spring tidal development. Debris were counted and classified across transects parallel to the coastline at progressively higher water marks. Plastics made up most of the AMD across all sites. More AMD was retained in the urban sites, consistent with their larger resident population density. Diversity of debris forms were consistent with the type of land use and population livelihood in each area. The greatest differences in abundance, diversity, and evenness were recorded between the lower tidal zones and the remaining inner transects consistent with sorting towards the coastal edge in favour of plastic items. Overall, differences across transects and sites suggested: 1) the canopy and root structure within the main body of the mangrove efficiently retained debris with little sorting; and 2) debris deposited closer to the edge is increasingly sorted and lost to the water body in favour of smaller plastic items, for a constant wind field and irrespective of neap-spring phases. The findings show that mangrove areas are vulnerable to a constant build of potentially harmful debris with selective leakage and sorting of materials back to the water body closer to their coastal edges. For Penang Island, the study highlights the areas in need of attention and prioritization, lists the types of debris needing proper management, and will aid in the future monitoring, mitigation and/or rehabilitation of these sensitive ecosystems.

## 1.0 Introduction

Anthropogenic marine debris (AMD) is one of the most serious threats to the environment, economy, and human health. Issues concerning AMD are now recognized internationally, alongside other major global challenges facing the marine environment including loss of biodiversity, ocean acidification, and sea level rise. Marine debris includes all materials discarded into the sea, on the shore, or brought indirectly to the sea by rivers, sewage, storm water, waves, or winds (NOAA, 2018). The most common materials that make up AMD are plastics, glass, metal, paper, cloth, rubber, and wood (NOAA, 2018). Of these, plastic can form up to 95% of the waste that accumulate on the shorelines, sea surface, and seafloor (Galgani et al., 2015). Bags, fishing nets, as well as food and beverage containers are the most common plastic items and constitute more than 80% of litter stranded on beaches (Topcu et al., 2013; Thiel et al., 2013).

AMD can originate from land or ocean. Land-based debris is accumulated from a variety of sources; littering or dumping, storm water discharges, outflow from industries, poor waste management practices, and occasional extreme natural events (i.e. tsunamis and hurricanes) (Aguilera et al., 2016; Goto and Shibata, 2015; Green et al., 2015; Khordagui and Abu-Hilal, 1994; Rech et al., 2014). Ocean-based debris is the result of bad management practices of cargo ships, fishing vessels, and off-shore oil and gas platforms (Astudillo et al., 2009; Edyvane et al., 2004; Hinojosa and Thiel, 2009; Hong et al., 2014; Sheavly and Register, 2007; Watters et al., 2010). The rise in tourism, agriculture, aquaculture, fisheries, and industrial activities worldwide have aggravated this environmental problem (Newman et al., 2015). With rapid population growth and urbanization, annual waste generation is expected to increase by 70% from 2016 levels to 3.4 billion tonnes in 2050 (Worldbank, 2019), stressing the need for mitigation measures for AMD.

Previous studies on AMD accumulation in the marine environment have mostly focused on sandy beaches, which are recommended for marine debris monitoring (Moore, 2008; Cheshire et al., 2009; Browne et al., 2015). However, the findings and lessons learnt from beaches cannot be extrapolated to more valuable coastal vegetated ecosystems such as mangroves (Costanza et al. 2014). Beaches are more open and dynamic, where longshore drift and daily ebb tides losses result in less than efficient net rates of accumulation and variable stocks (Erikson and Burton, 2003). Conversely, mangrove ecosystems, are characteristically less dynamic and more efficient retainers and accumulators of AMD (Martin et al., 2019), and potentially more vulnerable to the effects of AMD.

Mangrove forests cover about 132,000 km^2^ of subtropical and tropical shores (Hamilton and Casey, 2016). These forests are an integral part of the coastal environment and its functions and services provide for the well-being of flora and fauna on multiple trophic levels. Their key roles include carbon sequestration (McLeod et al., 2011; Almahasheer et al., 2017), coastal protection, habitat for marine life (Spalding et al., 2010; Duarte et al., 2013), and serves as a stop-over point for migratory birds (Yeok et al., 2016). They maintain water quality and clarity, filter pollutants and nutrients which can lead to algal blooms, and trap sediments originating from land (NOAA, 2018). Mangroves are extremely productive ecosystems that provide numerous goods and services which are conservatively estimated to be worth USD 186 million each year in terms of contribution from fisheries, timber and plant products, coastal protection, and tourism (WWF, 2019).

Mangroves are known to efficiently retain AMD even at low densities (Cordeiro and Costa, 2010). Their tendency to accumulate AMD is exacerbated by the structure of their pneumatophores (Martin et al., 2019) which is analogous to a fishing net that retains objects as they pass through and embeds them within its structure or the muddy substrate. Environmental factors such as seasonality (i.e. rainfall) (de Araújo and Costa, 2007; Ivar do Sul and Costa, 2013), hydrology (i.e. waves, currents, tides, local wind) (Corbin and Singh, 1993; Nagelkerken et al., 2001; Silva-Iniguez and Fischer, 2003; Schlining et al., 2013), coastline geography and sediment characteristics (i.e. muddy, sandy; extent of foreshore preceding the mangroves) (Cunningham and Wilson, 2003; Debrot et al., 1999; Mordecai et al., 2011), system entry sources (i.e. rural, peri-urban, urban, aquaculture, agriculture, industry, residential, commercial) (Santos et al., 2005; Sánchez et al., 2013; Leite et al., 2014), and population density can also influence AMD accumulation in mangroves. The size, weight, and type of the debris (Cordeiro and Costa, 2010; Possatto et al., 2015; Ivar do Sul et al., 2014; Martin et al., 2019) have also been reported to influence their accumulation and distribution within mangroves. This propensity towards accumulation is its strength in the protection of the coastline but also its weakness in accumulating AMD, causing detrimental, long-term effects to the ecosystem. These blue carbon ecosystems are becoming increasingly threatened by AMD accumulation.

Adding to the environmental factors and latent characteristics of the debris and vegetation, is the issue of direct/illegal dumping. Direct/illegal dumping not only harms natural ecosystems but poses danger to its community. For example, repeated illegal dumping of flammable materials like plastics is said to be the cause of a series of mangrove forests fires in Navi Mumbai, India, killing many mangrove trees and releasing smoke and carcinogenic fumes which affected nearby residents (Singh, 2018). Debris dumped into mangroves can be from on-site or off site. On-site debris dumping typically occurs when occupants of habitations on the mangroves directly dump rubbish (i.e. kitchen waste) into the mangroves. Such a case has been reported for a mangrove swamp in Sao Vicente Estuary, Brazil, where solid residues were associated with illegal dumping by occupants of habitations along the riverbanks (Cordeiro and Costa, 2010). Off-site debris dumping, on the other hand, involves debris from outside the mangroves being transported to and dumped into the ecosystem.

Considering the forests’ social, economic, and ecological importance, the issue of marine debris accumulation in mangroves have been largely neglected (Debrot et al., 2013). The paucity of research of AMD accumulation in mangroves is apparent in that only a handful of studies have been reported with most of them focusing on microplastic in mangrove sediments (Barasarathi et al., 2011; Lima et al., 2014; Mohamed Nor and Obbard, 2014; Lourenço et al., 2017; Naji et al., 2017). In the meta-analyses of spatial and temporal patterns of stranded intertidal marine debris worldwide in which 104 published scientific papers were reviewed, only four studies were found to involve mangroves (Browne et al., 2015). A further review of these four studies revealed only one had been conducted under the mangrove canopy itself (Cordeiro and Costa, 2010) whilst the remaining were carried out in creeks and beaches (Ganespandian et al., 2011; Singare, 2012; Debrot et al., 2013). The study conducted within the canopy had concentrated on the different densities at high and low tides and did not include a diversity and evenness analyses (Cordeiro and Costa, 2010).

Little is known on the nature and dynamics of AMD accumulation in mangroves. The influence of different land uses and environments in relation to retention, filtering, and sorting properties in these forests, remains unknown. There is also a paucity of information on where AMD tend to accumulate and whether their accumulation is selective at different parts of the mangroves. This information is crucial for several reasons including for effective management of AMD in mangroves which is usually constrained by organization resources and workforce availability. To date only two recent studies, have addressed some of these concerns (Ivar do Sul et al., 2014; Martin et al., 2019). The study by Ivar do Sul et al. (2014) was limited in the choice of AMD. They focused on the retention and exportation of a selected number of plastics forms deliberately released in mangrove habitats. The study by Martin et al. (2019), on the other hand, considered debris supplied from the open ocean. However, their study across the Red Sea supported a less than congested set of population centres, unlike regions such as Malaysia and the rest of Southeast Asia currently under pressure from AMD pollution. Nevertheless, both studies suggested that retention is a function of the mangrove and debris structures, but no indication of how this affected the overall retention or selection throughout the canopy.

To address this gap, first level assessments were carried out in Penang, a city-state located on the northwest coast of Peninsular Malaysia. Penang Island has 6.8 km^2^ mangroves left, much of which are constantly under threat from land use changes and marine pollution (Chee et al., 2017). The population relies on the many ecosystem services provided by mangroves through fisheries, wood harvesting, and tourism, to supplement the income of this state (Hamdan et al., 2012). The mangroves on the west coast of the island was also attributed for protecting the people and property during the 2004 Andaman tsunami (Alongi, 2008) that claimed over 250,000 lives worldwide—52 in Penang Island alone. With the increase in the pollution reports (Hezri et al., 2019) and frequency and severity of natural disasters due to climate change (Cheal et al., 2017), the city-state could substantially benefit from the findings of this study. Here we aim to characterize the different forms of debris across urban and peri-urban mangroves, constrained by any changes in wind field and neap tidal cycles. Sorting and retention of these forms was assessed from the changes in diversity, abundances, and evenness through the canopy, close to the coastal edge and into the main body of the mangrove stand.

## 2.0 Methodology

### 2.1 Study area

The study was conducted in two urban and two peri-urban mangroves on Penang Island (Fig. 1). These urban and peri-urban mangroves were categorized as such based on population density and land use, in a previous study (Nordhaus et al., 2019). Jelutong (JEL) and Free Trade Zone (FTZ) represented the urban sampling sites. Jelutong is located on the east coast, between a landfill and a fishing village, next to a highway. In terms of population livelihood and density at this site, there was some stratification to the north and south but generally, none between the edge and body of the mangroves. Free Trade Zone, on the other hand, is located on the southeast coast in an industrial zone. This sampling site is adjacent to Sungai Kluang river which is fringed by mangroves and stilted residences on the riverbank. There is no gradient in the population density and livelihood between the edge and the body of the mangroves, here.

**Fig. 1.**
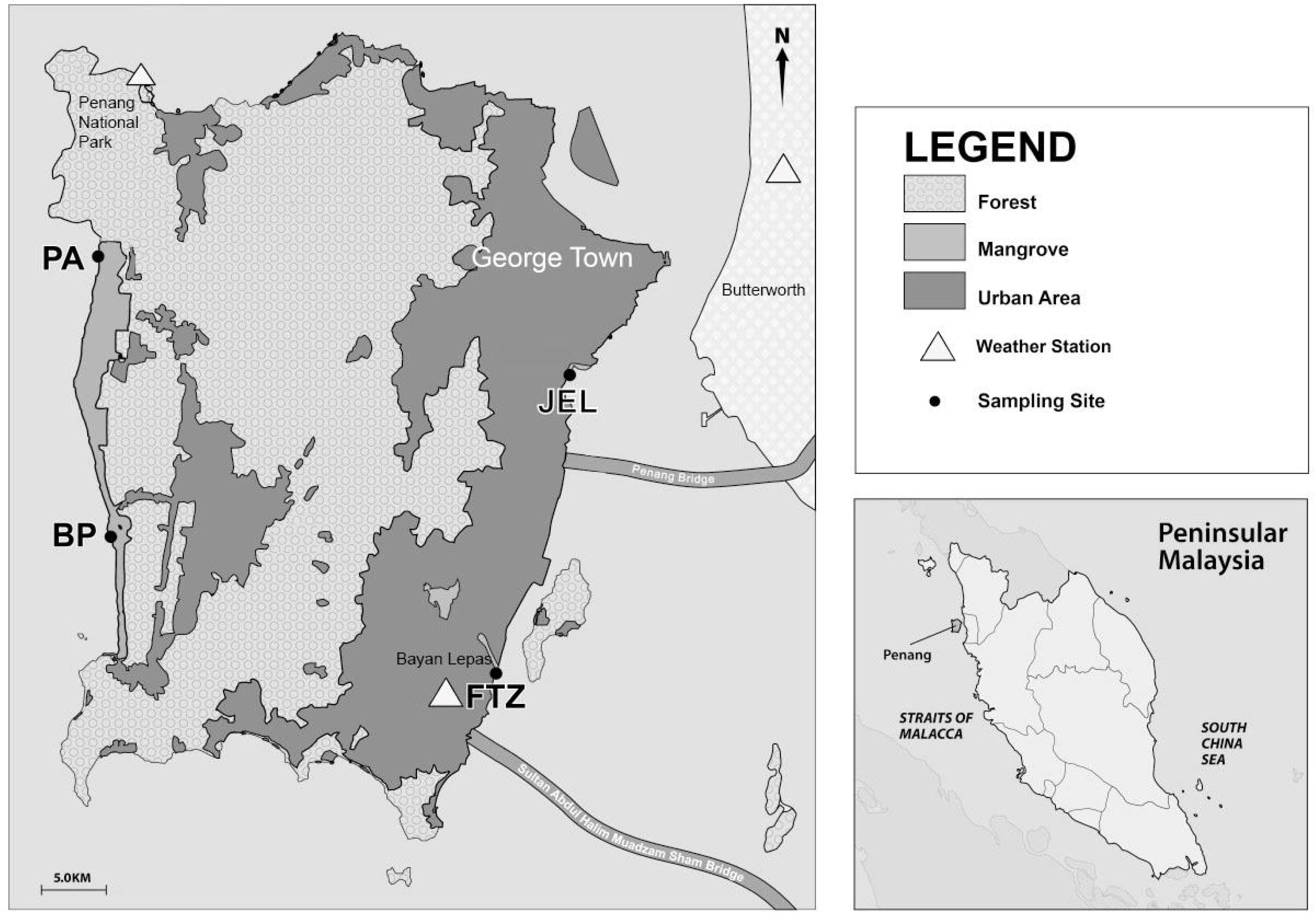
Sampling sites for anthropogenic marine debris accumulation in urban and peri-urban mangroves on Penang Island, Malaysia. Urban mangroves: Jelutong (JEL), Free Trade Zone (FTZ). Peri-urban mangroves: Balik Pulau (BP), Pantai Acheh (PA).

Pantai Acheh (PA) and Balik Pulau (BP) represented the peri-urban sites. Pantai Acheh is a coastal village on the north-west of Penang Island. It is the least urbanised and the most sparsely populated. There is a small fishing facility in the mangroves but, due to the remote location, there is far less human activity and disturbances in comparison to the other sites. Balik Pulau on the southwest coast has a slightly higher population and more urbanised. There is a mangrove forest reserve here, but large swathes of it are under threat to aquaculture and highway construction. Sampling sites and transect details are presented in Table 1.

**Table 1.**
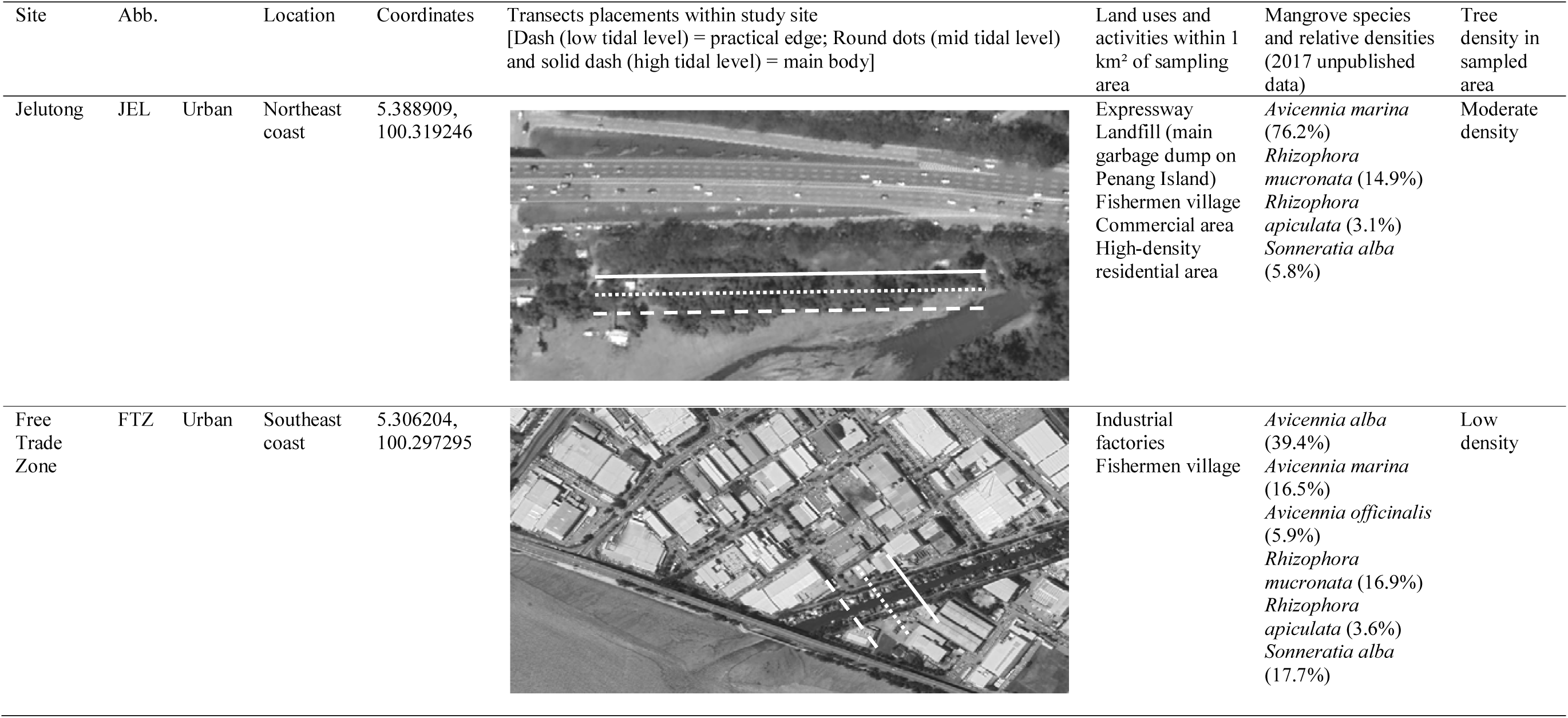
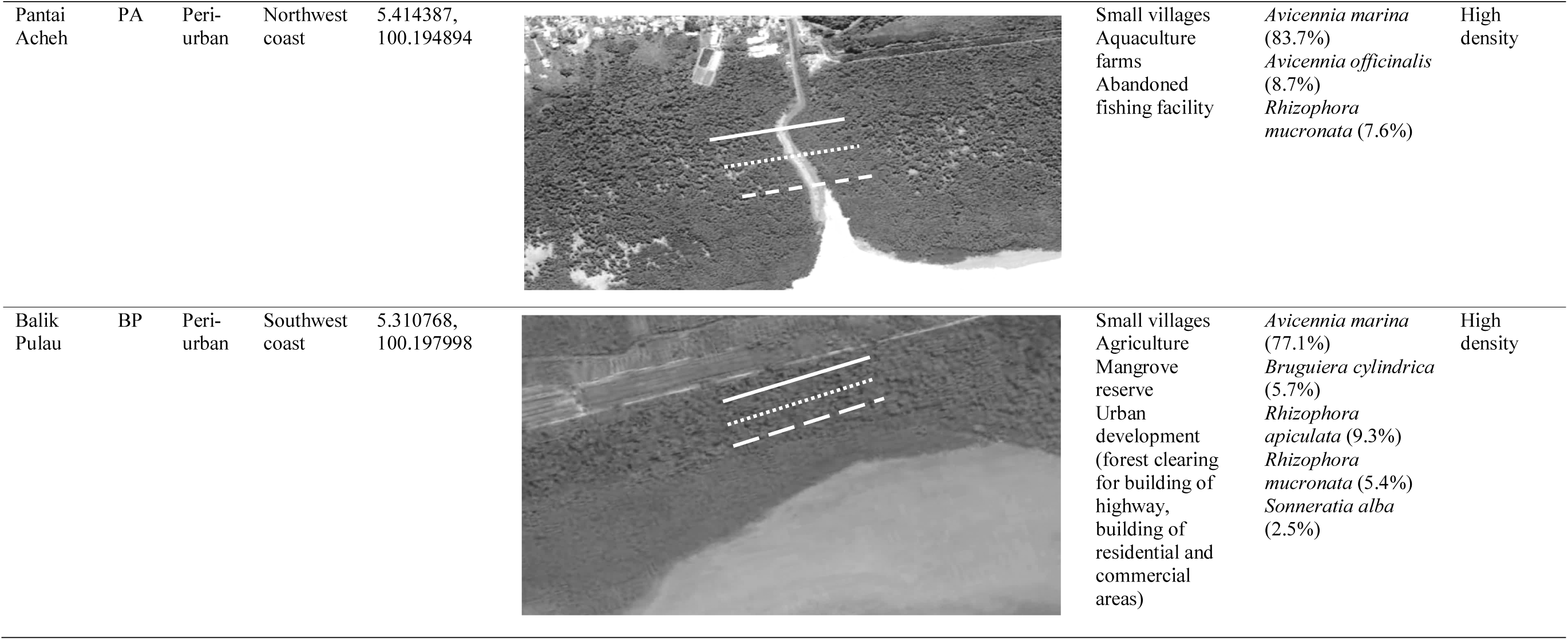
Sampling site description.

| Site | Abb. |  | Location | Coordinates | Transects placements within study site<br>[Dash (low tidal level) = practical edge; Round dots (mid tidal level) and solid dash (high tidal level) = main body] | Land uses and activities within 1 km<sup>2</sup> of sampling area | Mangrove species and relative densities (2017 unpublished data) | Tree density in sampled area |
| --- | --- | --- | --- | --- | --- | --- | --- | --- |
| Jelutong | JEL | Urban | Northeast coast | 5.388909, 100.319246 |  | Expressway<br>Landfill (main garbage dump on Penang Island)<br>Fishermen village<br>Commercial area<br>High-density residential area | *Avicennia marina* (76.2%)<br>*Rhizophora mucronata* (14.9%)<br>*Rhizophora apiculata* (3.1%)<br>*Sonneratia alba* (5.8%) | Moderate density |
| Free Trade Zone | FTZ | Urban | Southeast coast | 5.306204, 100.297295 |  | Industrial factories<br>Fishermen village | *Avicennia alba* (39.4%)<br>*Avicennia marina* (16.5%)<br>*Avicennia officinalis* (5.9%)<br>*Rhizophora mucronata* (16.9%)<br>*Rhizophora apiculata* (3.6%)<br>*Sonneratia alba* (17.7%) | Low density |
| Pantai Acheh | PA | Peri-urban | Northwest coast | 5.414387, 100.194894 |  | Small villages<br>Aquaculture farms<br>Abandoned fishing facility | *Avicennia marina* (83.7%)<br>*Avicennia officinalis* (8.7%)<br>*Rhizophora mucronata* (7.6%) | High density |
| Balik Pulau | BP | Peri-urban | Southwest coast | 5.310768, 100.197998 |  | Small villages<br>Agriculture<br>Mangrove reserve<br>Urban development (forest clearing for building of highway, building of residential and commercial areas) | *Avicennia marina* (77.1%)<br>*Bruguiera cylindrica* (5.7%)<br>*Rhizophora apiculata* (9.3%)<br>*Rhizophora mucronata* (5.4%)<br>*Sonneratia alba* (2.5%) | High density |

### 2.2. Anthropogenic marine debris (AMD)

Assessments were carried out between October and November 2018 over the monsoon transition period. For all four sites, three transects were placed parallel to the coastal edge (Table 1). The transect at the low tidal level ran as close as possible to the edge of the sea (henceforth referred to as the practical edge) and the other two transects were placed in the main body of the mangroves at mid and high tidal levels. Assessments of the total number and types of AMD for three 10 x 10 m (100 m^2^) quadrats were completed on each transect, totalling in 9 quadrats per sampling site. For more effective data collection, these quadrats were divided into one hundred 1 m^2^ sub-quadrats.

### 2.3 Environmental factors

Projected daily tidal ranges for September to December 2018 were obtained from tide tables (MetMalaysia, 2017). Hourly wind speed and direction data for the same period were obtained from three weather stations: WMKB (5.4659°N, 100.3912°E), WMKP (5.2971°N, Brassinosteroids (BRs) are a group of growth-promoting plant hormones, isolated and characterised from can 100.2769°E), and a research station in the Penang National Forest Reserve (5.49°N, 100.2025°E). The wind rose was plotted using the “openair” package in R (Carslaw and Ropkins, 2012).

### 2.4 Data analyses

Mean, standard deviation, and alpha diversity indices were calculated based on untransformed abundance data for each sampling site. Alpha diversity indices included: (1) Shannon-Wiener diversity index (H), the measure of diversity based on species richness and their relative frequency; and (2) Species evenness (EH), the measure of heterogeneity of a community based on distribution of relative frequency of species. A high H index would imply a diverse community structure while a high EH index would indicate an equally distributed community.

To address the similarity in terms of AMD community among the urban and peri-urban sampling sites, hierarchical cluster analysis was used to identify grouping of sites in accordance to multivariate community composition. Clusters were identified using the unweighted pair-group method with arithmetic mean (UPGMA) based on Chord similarity index. At the same time, analysis of similarity (ANOSIM) was used to statistically test whether there was a significant difference between all four sampling sites. If the groups of sampling sites were different in their community composition (AMD type), the compositional dissimilarities between groups would be higher than that within groups. Hence, ANOSIM R is based on the difference of mean ranks between groups and within groups. When a significant difference was detected, pair-wise post-hoc test was used to examine the correlation between groups. A large positive R value (-1 to +1) would signify dissimilarity between groups whereas 0 would indicate random grouping.

To address potential tidal fluctuation effect on AMD retention, we performed two-way ANOSIM test on each tidal level (low, mid and high) crossed between each sampling site. In addition, Similarity Percentage (SIMPER) with Chord similarity measure was performed to assess which taxa was primarily responsible for the difference between each tidal level within groups. All statistics were performed using Paleontological Statistics software package (PAST) version 3.22. A tidal curve was also generated using average daily means and subsequently smoothed with a 2-week moving average to illustrate variance over the spring-neap cycle in PAST. Raw data was submitted to Mendeley Data (Chee et al., 2019).

## 3.0 Results

### 3.1 Physical parameters

Throughout the sampling period, wind fields at the three weather stations did not vary before, during, and after the sampling period (September-December 2018). Wind fields were dominant at the northwest quadrant in Butterworth, southwest quadrant in Bayan Lepas, and south quadrants in Penang National Park (Fig. 6). Urban sites were sampled during a rising neap-spring tide whilst peri-urban sites were sampled during a falling neap-spring tide.

### 3.2 Site specific anthropogenic marine debris

#### 3.2.1 Abundance

The mangrove stands with the greatest AMD abundance were found along transects within the densely populated urban areas on the east coast of Penang Island. In JEL, which is located to the north and next to the island’s main landfill, the highest abundance was recorded at a total of 7,312 items (Table 2). Of these, 92.5% were plastic materials while the remaining 7.5% were non-plastics (Fig. 2a). The plastic debris ostensibly comprised of plastic bags as the largest fraction (2,046; 30.3%), followed by plastic sheets (1,343; 19.9%) and cutlery (995; 14.7%) (Fig. 3a). Non-plastic materials consisted mainly of glass and ceramic (261; 47.5%), followed by cloth (161; 29.3%) and rubber (81; 14.8%). The other urban site in the south, FTZ, which is located within the island’s industrial zone, 1,258 items were recorded. Of these, 77.7% were plastics (Fig. 2b). Like JEL, plastics bags (254; 30.0%) contributed the highest number of plastics followed by plastic sheets (248; 25.4%) and cutlery (117; 12.0%) (Fig. 3b) with non-plastic items following a similar hierarchy (Fig. 4b) of glass and ceramic (98; 34.9%), rubber (69; 24.6%) and cloth (54; 19.2%). Other plastics items found within the urban mangroves include bottles, bottle lids, food containers, fishing nets, lighters, cigarette butts, mesh bags, foam, and fragments. Non-plastic items like paper, cardboard, and metal were also spotted.

**Fig. 2.**
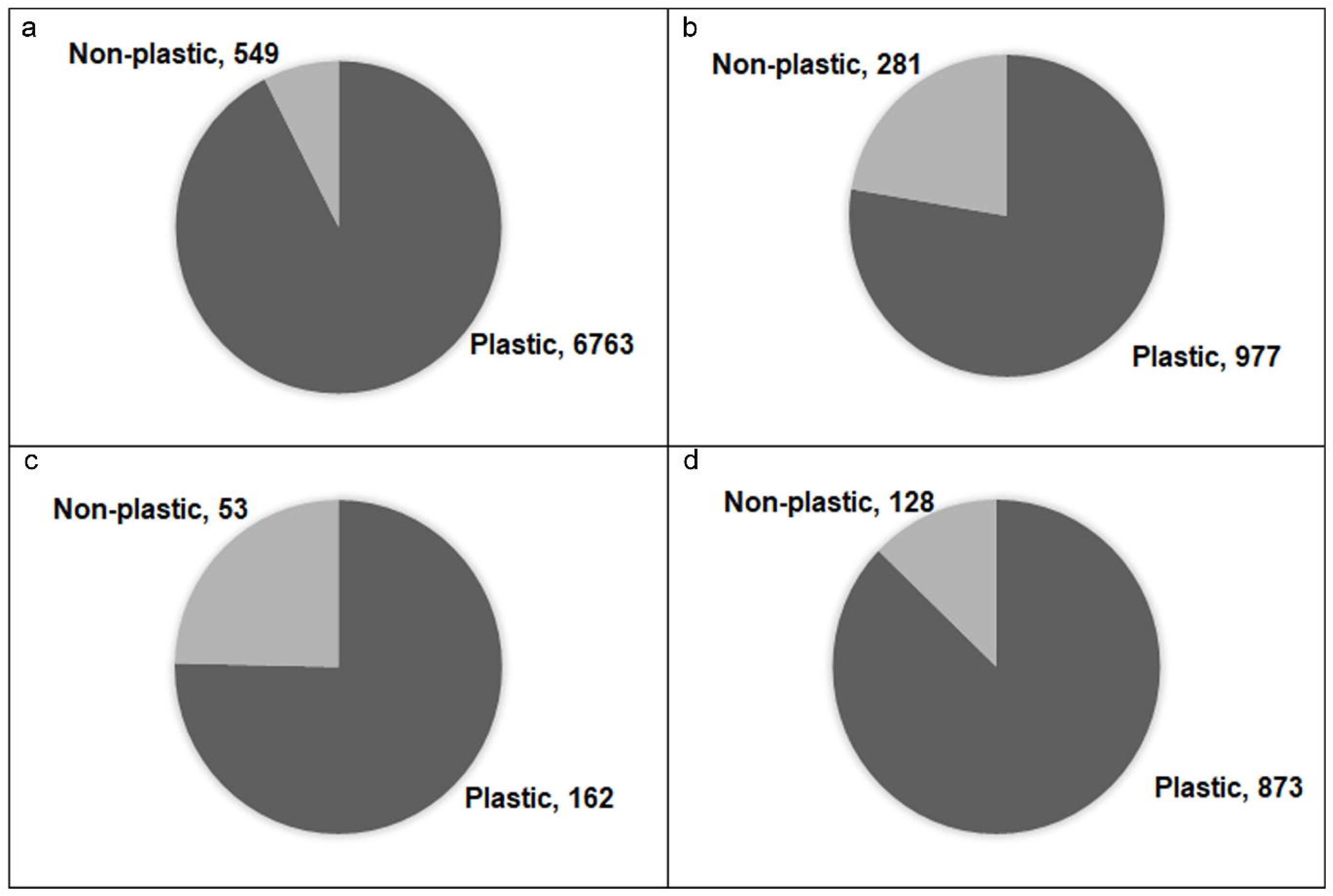
Proportions of plastic versus non-plastic debris accumulated at (a) Jelutong (JEL), (b) Free Trade Zone (FTZ), (c) Pantai Acheh (PA), and (d) Balik Pulau (BP).

**Fig. 3.**
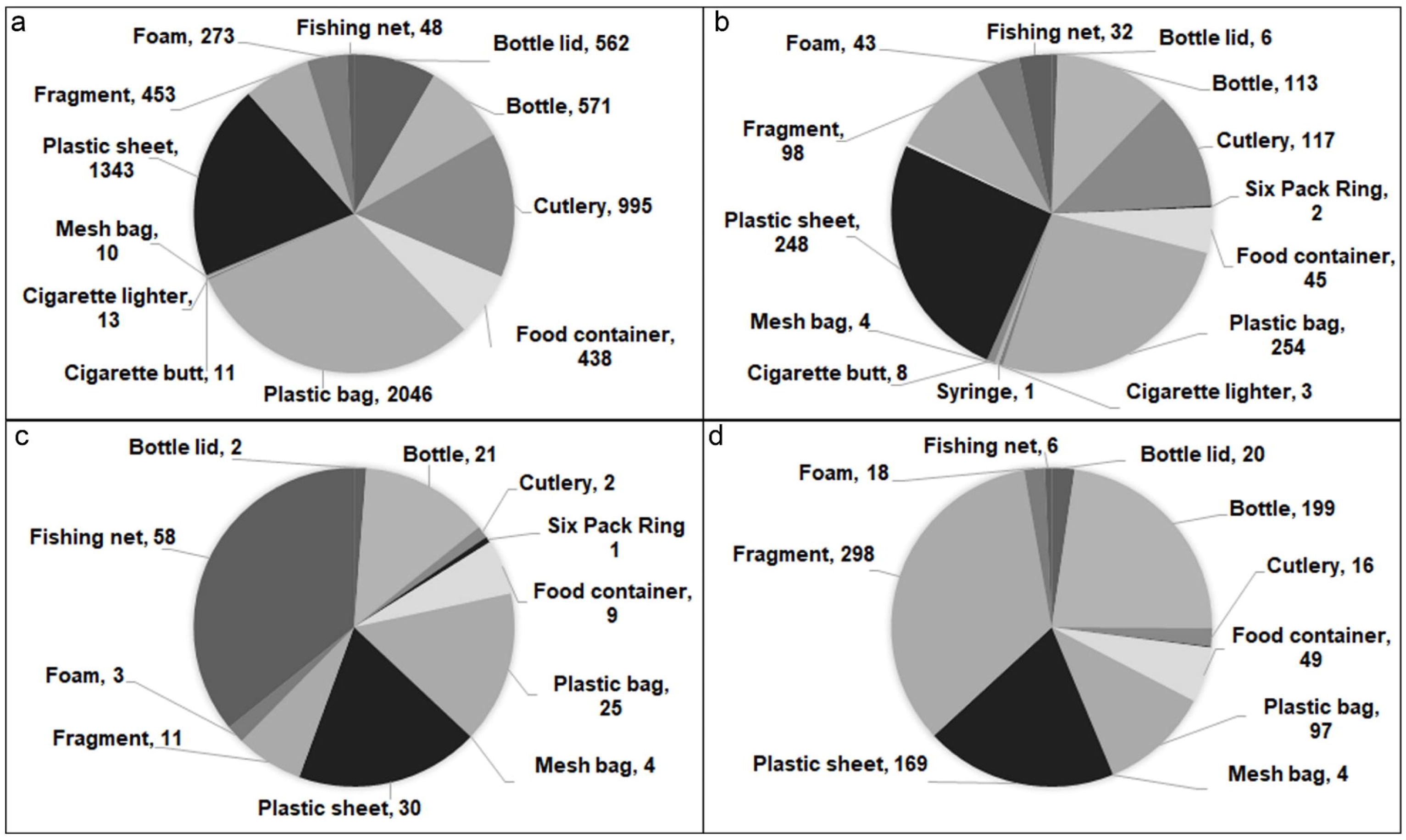
Proportions of plastic debris accumulated at (a) Jelutong (JEL), (b) Free Trade Zone (FTZ), (c) Pantai Acheh (PA), and (d) Balik Pulau (BP).

**Fig. 4.**
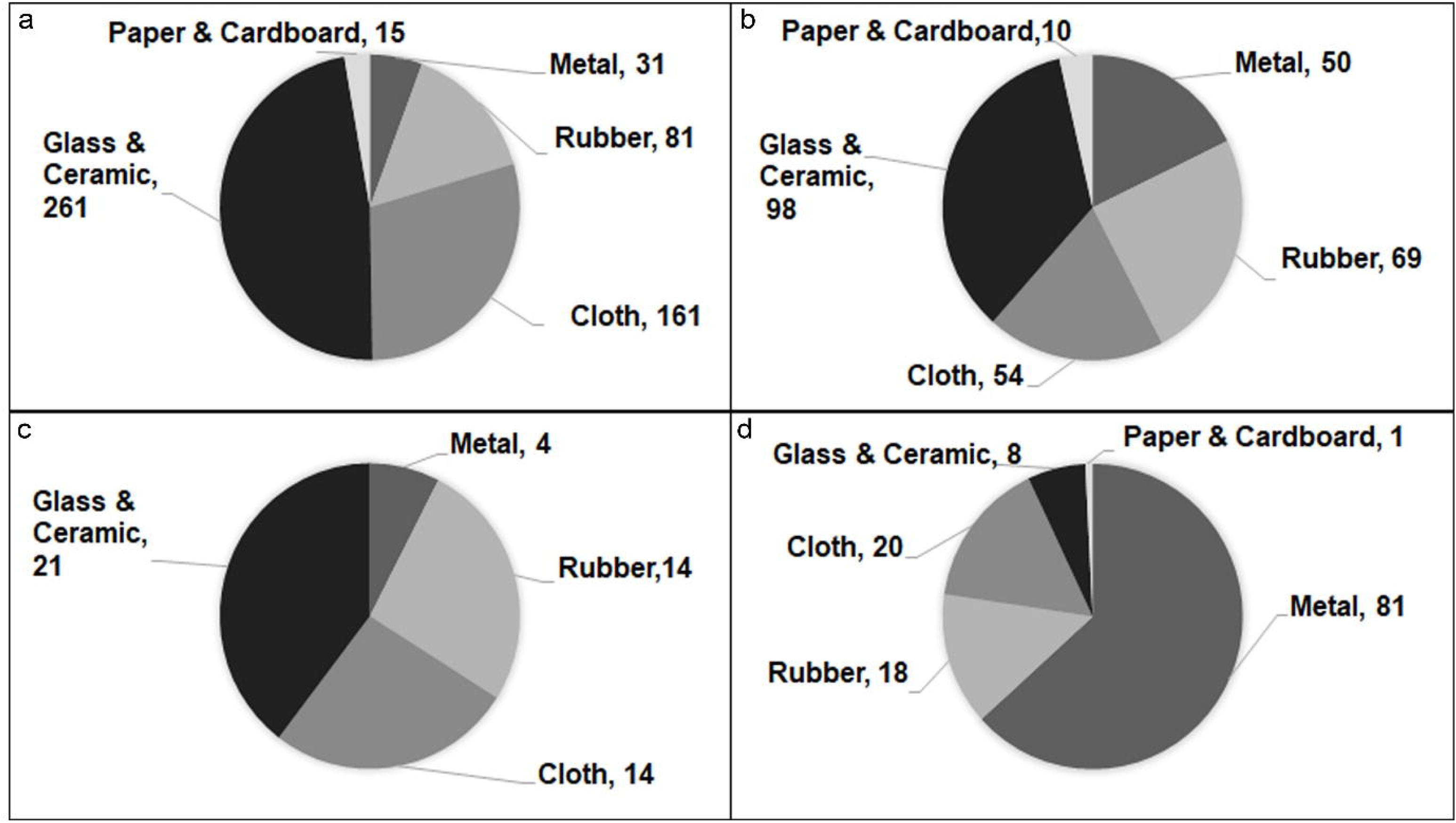
Proportions of non-plastic debris accumulated at (a) Jelutong (JEL), (b) Free Trade Zone (FTZ), (c) Pantai Acheh (PA) and (d) Balik Pulau (BP).

**Table 2.**
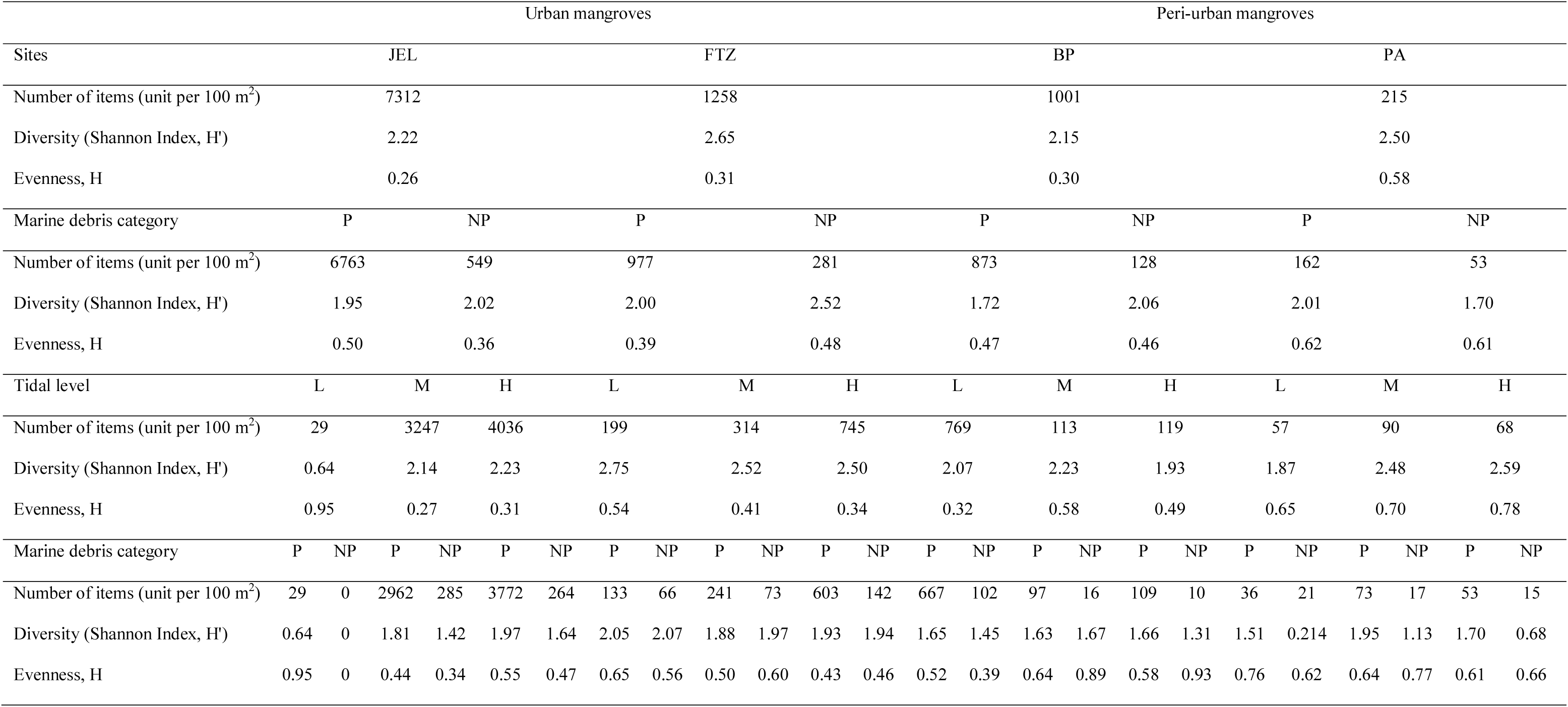
Number, diversity, and evennes of anthropogenic marine debris in each sampling site and stratification according to tidal levels.

The sparsely populated peri-urban sites on the west coast had notably lower abundances of AMD. Located at an abandoned fishing facility to the north of the island, PA recorded the least abundance of AMD of all the four peri-urban and urban sites, with 215 items. Of these, 75.3% were plastics (Fig. 2c). In contrast to the urban sites, within PA, fishing nets (58; 35.8%) were the most common plastic items found followed by sheets (30; 18.5%) and plastic bags (25; 15.4%) (Fig. 3c) but with a similar non-plastic hierarchy (Fig. 4c) of glass and ceramic (21; 39.6%), rubber (14; 26.4%), and cloth (14; 26.4%). The other peri-urban site to the south of the island, BP, which is located within a mangrove reserve, recorded 1,001 items. Plastic items constituted most of the AMD items (87.2%) (Fig. 2d). Unlike the other sites, plastic fragments dominated (298; 34.1%) followed by bottles (199; 22.8%) and sheets (169; 19.4%) for non-plastics, with metals emerging as dominant (81; 63.3%), followed by cloth (20; 15.6%), and rubber (18; 14.1%) (Fig. 4d). Other plastic items found in the peri-urban mangroves were plastic bags, food containers, cutlery, bottle lids, six pack rings, fishing nets, and foam. Non-plastic items such as paper and cardboard were found in BP but not in PA.

#### 3.2.2 Diversity

The diversity of AMD was found to be similar among all sites (Table 2). The highest diversity (H’ = 2.649) was recorded in FTZ whilst the lowest diversity (H’ = 2.149) was recorded in BP. The evenness was similar among JEL (H = 0.2629), FTZ (H = 0.3143) and BP (H = 0.2958) but significantly lower in PA (H = 0.577). Nevertheless, despite the similarity, a significant difference was detected in the AMD “community” among sampling sites (p = 0.0003, R = 0.5864, Table 3).

**Table 3.** ANOSIM pairwise table (post-hoc R-values). Values indicate degree of similarity between accumulated marine debris in peri-urban (BP = Balik Pulau, PA = Pantai Acheh) and urban sites (FTZ = Free Trade Zone, JEL = Jelutong). * indicate significant values (p <0.05).

|  | FTZ | JEL | BP | PA |
| --- | --- | --- | --- | --- |
| FTZ |  | -0.04 | 0.78* | 0.52* |
| JEL | -0.04 |  | 1 | 0.70* |
| BP | 0.78* | 1 |  | 0.74 |
| PA | 0.52* | 0.70* | 0.74 |  |

Correspondingly, the HCA dendrogram revealed three clusters: 1) PA (peri-urban), 2) BP (peri-urban), and 3) JEL and FTZ (both urban) (Fig 5). SIMPER analysis showed that dissimilarity between the urban sites, FTZ and JEL, was low (11%) and confined to differences in relatively high numbers of bottle caps and ceramic construction material (a cumulative 44%) across the mangrove stands. Comparisons between the rest of the sites showed differences in AMD “community” with the largest dissimilarity (91%) between BP and PA contributed by fishing nets in PA and fibreglass fragments in BP. In total, these two types of debris cumulatively accounted for 76% dissimilarity between the two sites. The percentage of dissimilarity was similar between JEL and BP (77%) and JEL and PA (78%). Between JEL and BP, the differences were mainly contributed by fibreglass fragments and plastic bags in JEL (71%), whilst between JEL and PA the differences were largely contributed by plastic bags and cutlery in JEL (78%). Percentages of differences were lower between FTZ and BP (58%) and FTZ and PA (58%). Fibreglass fragments in BP and plastic bags in FTZ contributed most of the dissimilarity (65%) between FTZ and BP, whilst fishing nets in PA contributed55% of the dissimilarity between FTZ and PA.

**Fig. 5.**
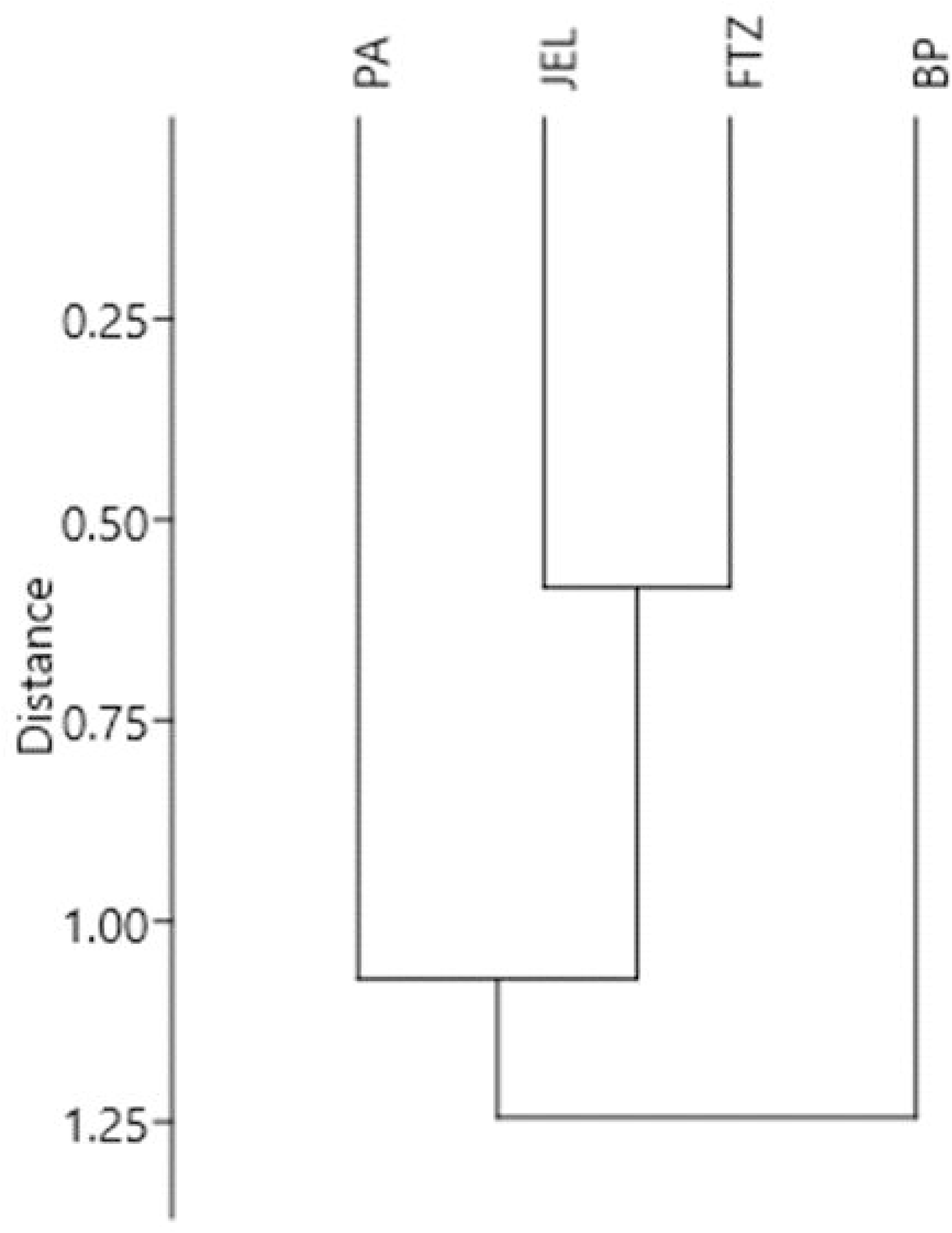
Dendrogram of relationship between urban (JEL = Jelutong and FTZ = Free Trade Zone) and peri-urban (PA = Pantai Acheh and BP = Balik Pulau) sites.

### 3.3 Anthropogenic marine debris variability within sites

On a rising neap-spring tide (Fig. 7), patterns of AMD accumulation at different tidal levels were similar in the two urban mangroves. Abundances of AMD in JEL and FTZ were lower at the practical edges than in the main body of the mangroves (Fig. 8). Although not significantly different, the differences in the abundance of AMD accumulated at the practical edge versus the main body was more pronounced in JEL than in FTZ. In JEL, closest to the edge, showed the greatest transition in AMD abundance from the practical edge to the main body. The practical edge here recorded the lower diversity (H’ = 0.64), whilst the main body recorded similar and higher diversities (H’ = 2.14 to 2.24). Evenness was higher at the practical edge (H = 0.95) compared to the main body (H = 0.26 to 0.31). In FTZ, which was slightly further from the edge, the transition was less pronounced. Similar diversities were observed throughout the tidal levels ranging from H’ = 2.50 to 2.52 in the main body to H’ = 2.75 at the practical edge. Evenness here was higher at the practical edge (H = 0.54) compared to the main body (H = 0.34 to 0.41). While the abundances were different within the main bodies of the urban sites, the diversity and evenness were similar. However, in the case of the practical edges, differences in diversity were detected. SIMPER analyses revealed that this was mainly attributed by the accumulation of plastic cutlery, sheets, bottles, and bags in the main body of JEL, and the retention of plastic cutlery at the practical edge as well as plastic bottles, sheets, construction material, and fibreglass fragments in the main body of FTZ (Table S1).

**Fig. 6.**
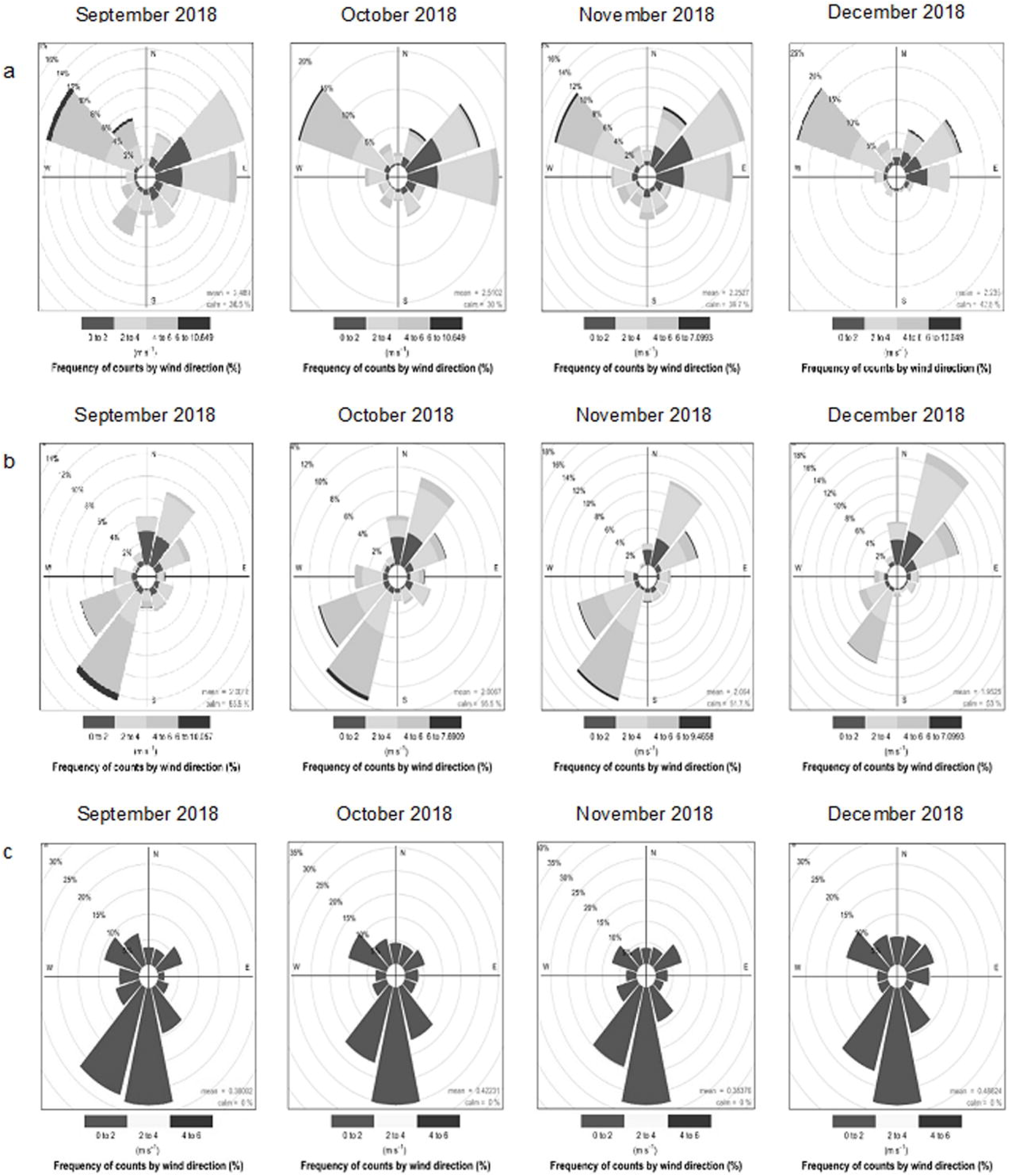
Wind rose plot for the months of September, October, November and December 2018 retrieved form weather stations in (a) Butterworth, (b) Bayan Lepas and (c) Penang National Park.

**Fig. 7.**
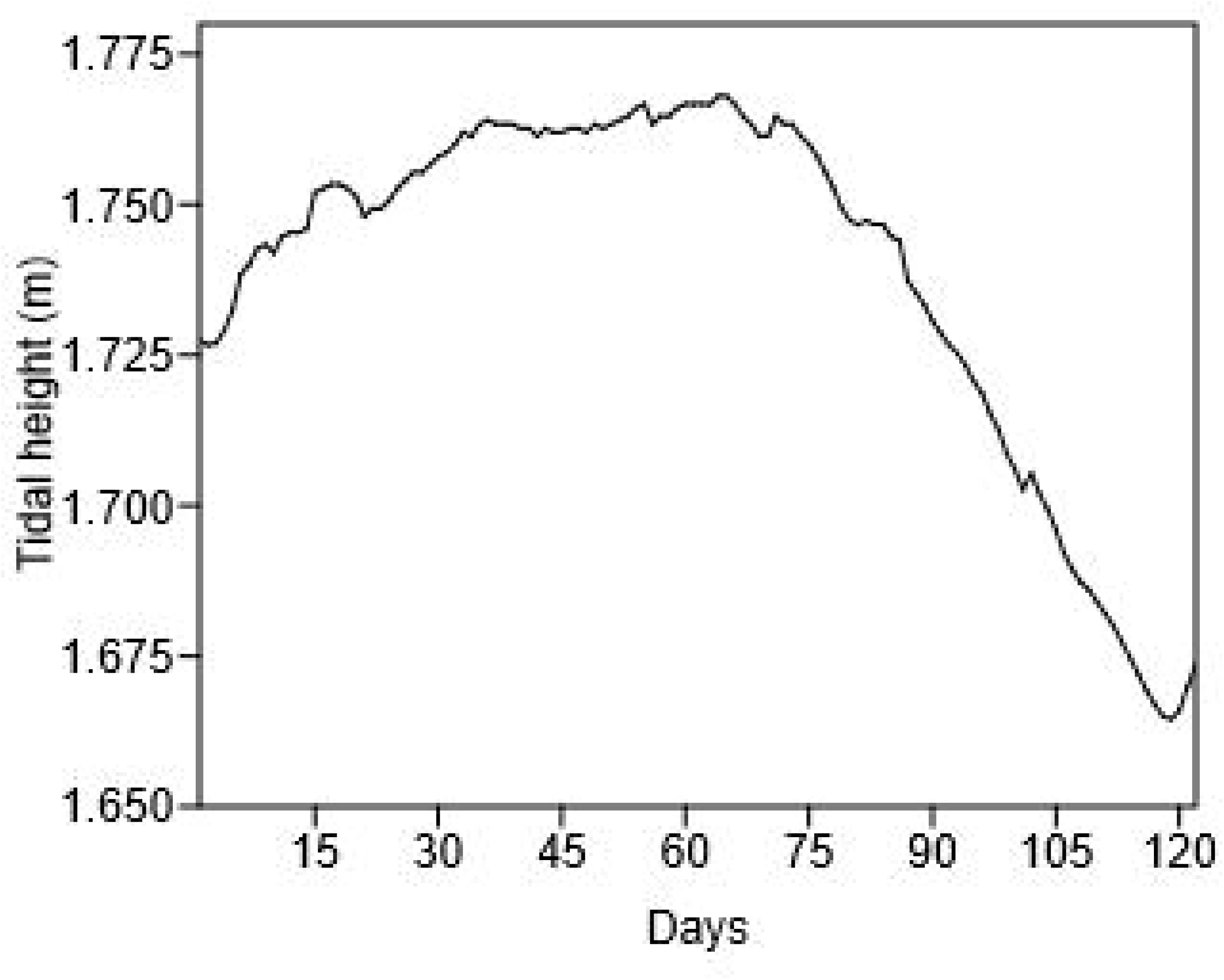
Monthly tidal range from September to December 2018 smoothed by moving average at 14 points (α = 0.50).

**Fig. 8.**
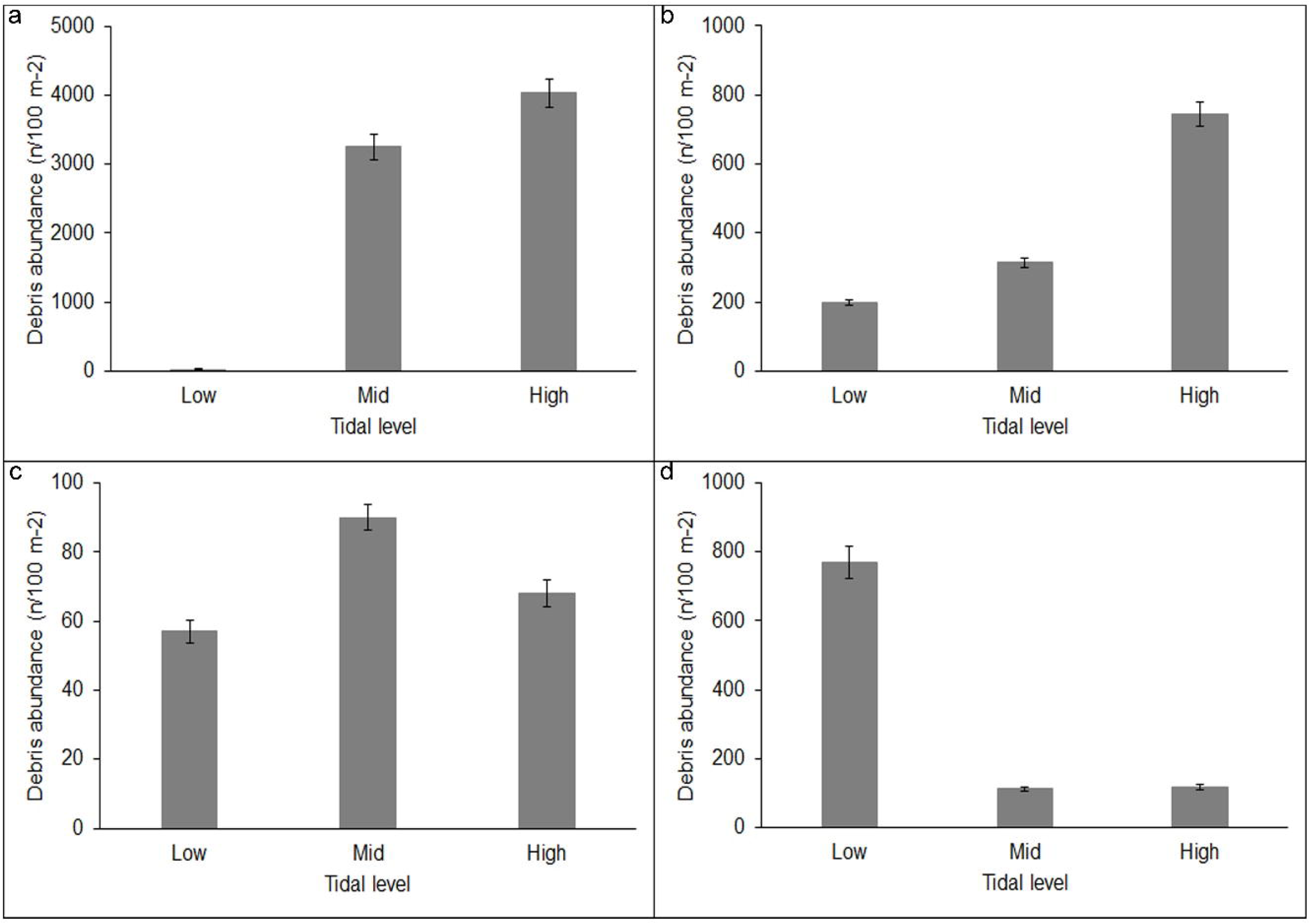
Abundance of anthropogenic marine debris accumulated at different tidal levels at (a) Jelutong (JEL), (b) Free Trade Zone (FTZ), (c) Pantai Acheh (PA), and (d) Balik Pulau (BP).

The peri-urban sites, which were both sampled on a falling neap-spring tide, displayed either decreasing or variant patterns in terms of AMD accumulation. The practical edges in these two sites were further inland compared to the practical edges in the urban sites. In contrast to the urban sites, BP displayed a decreasing pattern. Debris abundance was higher at the practical edge and lower through the main body. Although the abundances were different, the diversity and evenness remained similar. Diversity ranged from H’ = 2.06 at the practical edge to H’ = 1.93 to 2.23 in the body of the mangroves while evenness ranged from H = 0.32 at the practical edge to H = 0.49 to 0.58 in the body of the mangroves. The differences in diversity between the practical edge and the main body was mainly caused by the accumulation of fibreglass and plastic bottles at the practical edge (Table S1).

Inconsistent with the rest of the sites, PA displayed a variant pattern with higher AMD abundance in the mid intertidal zone compared to the practical edge and high intertidal zone. The practical edge had a slightly lower diversity (H’ = 1.855) compared to the main body of the mangroves (H’ = 2.475 to 2.588). Evenness was slightly elevated in the high intertidal zone (H = 0.7824) compared to the practical edge (H = 0.6454) and the mid intertidal zones (H = 0.6993). The differences in diversity between the practical edge and the main body in PA was almost wholly induced by the retention of construction material seemingly directly dumped at the practical edge, as well as fishing nets and bottles in the main body of the mangroves (Table S1).

To eliminate the confounding factor of direct dumping of dominant AMD at the two peri-urban sites, additional analyses were performed with these items removed. After removing construction material from BP and PA, as well as fishing nets from PA, we found that there were no differences in abundance, diversity and evenness.

## 4.0 Discussion

### 4.1 Anthropogenic marine debris composition

The dominance of plastic items observed at all sites supports the plethora of global studies that have quantitatively described marine debris from beaches (Frost and Cullen, 1997; Derraik, 2002; Ivar do Sul and Costa, 2007; Sheavly and Register, 2007; Moore, 2008, Ryan et al., 2009; Leite et al., 2014; Possatto et al., 2015) and mangroves (Sivasothi, 2002; ICCS, 2009). This recurring pattern of plastic contamination reflects its versatility and subsequent wide-scale high use by modern society, variability, low-density (i.e. propensity to float), and resilience to environmental degradation (Derraik, 2002; Katsanevakis and Katsarou, 2004).

The abundances of different AMD classes are consistent with the dominant human activity at each site (Abu-Hilal and Al-Najjar, 2004). Large proportions of plastic bags, food containers, bottles, and cutlery were found in urban sites, while fishing nets, plastic sheets, bags, and bottles were found in greater magnitudes in peri-urban sites, because of their environmental persistence, ample utilization, and low reuse and recycling rates, as has been observed in other regions (Derraik, 2002; Santos et al., 2005; Cordeiro and Costa, 2010). In particular, JEL, located adjacent to the Jelutong Landfill (the main garbage dump on Penang Island), had almost six times more AMD compared to FTZ. The landfill had recently been reported to be operating over its capacity and potentially causing harm to the environment (Mok, 2019). In FTZ, located in an industrial zone with round-the-clock human mobility, occurrences of fibreglass, construction material, and cutlery were the highest. In the peri-urban sites, PA and BP, where fishing was the dominant human activity, more fishing nets and fibreglass fragments (the main material used to build fishing boats), were found.

A positive relationship between the population density and abundance of AMD was observed. The urban mangroves (JEL and FTZ), located on the east coast of Penang Island where most of the island’s development (see Fig. 4 in Chee et al., 2017) and population is concentrated, recorded significantly more AMD compared to the peri-urban mangroves (BP and PA) sampled in this study. The relationship between population size and coastal contamination has previously been recorded in several studies (Barnes, 2005; McGranahan et al., 2007; Browne et al., 2011, Browne et al., 2015). Pollution in intertidal areas has been directly linked to the number of inhabitants in its vicinity, leading to largely populated areas having greater amounts of AMD as opposed to less populated areas (Browne et al., 2011; Seto, 2011; Yonkos et al., 2014). In this study, population size was potentially the driving factor for higher abundances of AMD in the urban sites, JEL and FTZ, and the elevated number of AMD in BP compared to PA. For the latter, even though both areas are forged from an interaction of urban and rural land use, BP located on the south-west of Penang Island has a considerably higher population density compared to PA on the north-west which consists of a small fishing village.

Aside from population density, other factors such as tree density and the inferred differential ability to retain AMD maybe part of the equation. This presumption can be inferred from the abundances of AMD accumulated in the two urban sites. JEL, which had moderate tree density, accumulated higher abundances of AMD compared to FTZ, which had low tree density (Table 1). It is presumed that mangrove roots can also trap objects (like floating plastic) transported by currents but to date, there is no indication how this affects the overall retention or selection throughout the canopy (Martin et al., 2019). In this study, tree stands and roots were observed to be thinning towards the coastal edge at the urban site, JEL, where the practical edge was at the coastal edge. Coincidentally, AMD was also found to be least abundant at the coastal edge and the number of plastic bags, bottles, cutlery, and food packaging were markedly reduced compared to the main body. Further investigations need to be carried out to test this phenomenon.

Eriksson et al. (2013) and Smith and Markic (2013), found that the loss and variance of open beach debris appeared to be driven at the scale of semi-diurnal to diurnal falling tides. Sadri and Thompson (2014), also found that this was also the case for the more mobile micro- and macroplastics which also show considerable variation on beaches at small timescales within a day. Therefore, it was initially surprising that we found no evidence for confounding of this retention dynamic at different monthly rising and falling spring-neap cycles. The patterns of abundance, diversity, evenness, and AMD accumulation at each site were either similar or did not require this explanation to describe the variance. The spring-neap relative invariance may well reflect the mangrove systems’ less dynamic character over beaches in its greater ability to retain material. In other words, the current mangrove stock is a result of a slow accumulation of material and not a complex beach dynamic balance that depends on the state of the tide and previous accumulated state. Other than tide, it has also been recognised that the rate of supply for net accumulation and sources of AMD is also dependent on the wind field. However, for this study, any differences between sites the result of sampling at different times was not confounded as it was also observed that these wind fields had remained in the same quarters for each site before and throughout the sampling period.

### 4.2 Anthropogenic marine debris dynamics

Interestingly, an edge effect was observed in the urban sites. Abundances were lower but diversities and evenness were higher at the practical edges compared to the main bodies of these mangroves. This is consistent with both sorting and less retention by diurnal tidal movements for which effects are more prominent closer to the coastal edge where tree densities are markedly lower than in the main body. Lower tree densities are presumed to result in less impediments to the free circulation of floating residues (Cordeiro and Costa, 2010). The edge effect was more prominent in FTZ than JEL in which sorting appeared to retain plastic cutlery over other forms of debris at its practical edge. The effect, however, did not appear to extend through the main body of the mangrove stand. There were higher AMD abundances that showed similar diversities and evenness consistent with relatively unimpeded transport into the mangroves on a rising tide as initially suggested by Cordeiro and Costa (2010), and efficiently retained after deposition. Thus, it appears that not only sorting may be happening towards the edge but there is a general fall in abundance and retention. The reasons behind sorting and falling retention are unclear. Changes in retention may stem from a change in hydrodynamic parameters, as arguably determined by the mix and density of mangrove structures such as pneumatophores and trunk as well as mud. Generally, pneumatophores are more abundant and longer in deeper water, often represented by the low tidal mark, and more recently seen to be associated with different forms of debris, such as plastic bags and nets (Martin et al., 2019). It is thus, not inconceivable that sorting as a selective retention of the smaller plastic components, as exemplified at JEL, may be the result of changing mangrove structure. Alternatively, the narrow width of the mangroves observed at both urban sites might also allow AMD to be pushed further and higher with each rising tide, with lighter and more buoyant debris (plastic bottles, sheets, and cutlery) having a greater tendency to accumulate there. Indeed, higher grounds have been reported to retain significantly more plastic items (Ivar do Sul et al., 2014) with debris most highly concentrated at natural wrack lines (Viehman et al., 2011). Clearly, these speculations require more detailed and extensive measurements of pneumatophores and trunk densities and sizes, not forgetting the role of muddy bottoms for the smaller components.

The edge effect was not as apparent in the peri-urban sites. We suggest that this is a function of practical sampling accessibility, where the practical coastal edges were further inland compared to those in our urban sites. Here, variability was observed throughout the main bodies of the two peri-urban mangroves. Nevertheless, higher abundance was observed at the practical edge in BP. Evenness and diversity were the same at all tidal levels indicating no selection or sorting. The main difference between BP and the other peri-urban site, PA, was the fact that even though the practical edge in BP was not close to the coastal edge, they were at the edge of naturally recruiting mangroves. Prominent differences in densities and arrangements of mangroves stands in the mangrove reserve versus the naturally occurring mangroves on the seaward side were observed during sampling. In fact, difficulty in accessibility into the naturally occurring mangroves was the reason for the placement of the seaward transect at this practical edge. Therefore, although there may not be an edge effect between the planted mangroves in the reserve and the sea, there may be an edge effect between planted and naturally occurring mangrove trees at this site. The naturally occurring mangroves were noticeably denser and is likely the reason for more AMD accumulating at the practical edge, but the nature of this retention is not selective for specific kinds of debris. Through the main body there was no difference in abundance indicating free movement and translocation of AMD through it. Contradictory to the urban sites, BP was sampled on a falling tide possibly exporting AMD seaward accumulating at the edge of the naturally occurring mangroves. Retention on a rising diurnal tide is also possible and subject to further studies.

Interestingly, while evidence of dumping was the most apparent at this peri-urban site, removing this variable did not significantly affect the diversity, evenness, and abundance. The greater driver here seems to be the continuous pressed vector of AMD transport into the mangroves from the sea and the ability of the mangroves to retain the AMD, over time. Clearly while dumping should be discouraged, resources directed to this relatively stochastic input would be effectively utilised towards managing debris at their sources and preventing them from entering the oceans.

It is interesting to note that monthly tidal regimes seem to have little influence of the abundance, diversity, evenness, and patterns of AMD accumulation at each site. On the contrary, diurnal tidal movements was observed to have a stronger influence on these parameters, aiding sorting in the narrower, urban mangroves, retention in denser mangroves, and intertidal movement in sparser mangroves. This is consistent with previous studies that concluded debris loads could vary considerably between days and accumulation rates estimated from monthly rather than daily collections are underestimated by up to an order of magnitude (Eriksson et al., 2013; Smith and Markic, 2013). Micro- and macroplastics can be highly mobile and show considerable variation at small timescales, even within one day (Sadri and Thompson, 2014). It was also observed that differences in the said parameters were not constrained by previous wind fields as these wind fields had remained in the same quarters before and throughout the sampling period.

### 4.3 Management recommendations

Although more stringent laws and enforcement can help curb the issue of direct dumping in mangroves, controlling this act alone will not significantly decrease the accumulation of AMD in mangroves. Management of debris at the source (e.g. plastic bags in urban mangroves, fishing equipment in peri-urban mangroves) through practices like reducing their generation, reusing, and recycling is recommended and would likely play a bigger role in preventing more debris from entering these ecosystems. Clean-ups can be organized to remove existing AMD, but these should not only concentrate on beaches but also in mangroves which have long been neglected and have greater abilities in retaining debris. Clean-ups in larger magnitudes and capacity is needed in urban mangroves where there are more contributors of debris compared to peri-urban mangroves. Removal of pollutants should be prioritised in mangrove bodies and denser mangroves where AMD accumulation is higher. The edges of the mangroves should be cleared more frequently compared to the main body to avoid debris from being lost to the water body after which, management would be increasingly difficult. Clean-up programs can be carried out throughout the year as monthly tidal regimes were observed to have little to no effect on the abundance, diversity, or evenness of AMD accumulated in mangroves. The key mechanism for minimizing AMD in peri-urban areas requires the integration between the local government and civil society, with the former responsible for regular bulk collection and disposal, and the latter for disposal at clearly identified locations. Standard operating procedures and management strategies should be reviewed and revamped periodically to account for changes in the nature of challenges and threats faced by mangroves.

Swift action is needed to avoid further accumulation of debris of the mangrove ecosystems. A recurring concern of such debris is the obvious visual pollution and economic repercussions for the tourist and marine industries associated with unwanted material either deposited in the mangroves or entangling and damaging equipment (e.g. Barnes et al., 2009; Derraik, 2002). At an ecosystem level, common items found in this study (i.e. plastic bags, sheets, fragments, fishing nets, glass, ceramics) can negatively influence biota via the absorption of harmful chemicals, transport of non-native marine species to new habitats on floating objects, crushing of vegetation or reduction of light levels needed for growth, causing injuries and death among marine animals, and bioaccumulating up trophic levels (Winston, 1982; Derraik; 2002; Teuten et al., 2009; Uhrin and Schellinger, 2011; Rochman et al., 2013; Vegter et al., 2014). There is also potential link between mangrove pollution and carbon sequestration, as stress from pollution can lead to mangrove mortality and less productive mangrove ecosystems (UN Environment, 2017). Public awareness, education, as well as advocacy campaigns can help change mindsets by stressing on the importance of conserving mangroves and its effect, otherwise.

## 5.0 Conclusion

One of the biggest regional threats to mangroves is improper management of waste. The ability of mangroves to retain land- and ocean-based debris harms the ecosystem, the species living there, and even human health. Anthropogenic marine debris accumulation in mangroves is understudied. Here, we compared abundance, diversity, evenness, and patterns of accumulation of AMD in urban and peri-urban mangroves around Penang Island, Malaysia. Like previous studies on AMD accumulation in the marine environment, plastics made up most of the AMD at all sites. More AMD was accumulated in urban compared to peri-urban sites, consistent with their larger resident population density. Highly diverse debris forms were consistent with land use and population livelihood in each area. At the practical edges within the lower tidal zones of the urban sites, we observed evidence of sorting in favour of plastic items. The greatest differences in abundance, diversity, and evenness were recorded between the lower tidal zones and the remaining inner transects consistent with sorting towards the coastal edge in favour of plastic items. These patterns of change and differences across transects and sites suggested: 1) the canopy and root structure within the main body of the mangrove efficiently retained debris with little sorting; and 2) debris deposited closer to the edge is increasingly sorted and lost to the water body in favour of smaller plastic items, for a constant wind field and irrespective of neap-spring phases. The findings highlight mangrove areas are vulnerable to a constant build-up of potentially harmful debris. They also stress the need to consider variability in abundance, diversity, and accumulation of materials across sites and within the canopy and root system to ascertain the nature of the sink and the leakage of materials back to the water body. This study presents areas in need of attention and prioritization in Penang, lists the types of debris needing proper management at the source, calls for swift action from the local government and civil society, and will aid in the future monitoring, mitigation and/or rehabilitation of these sensitive ecosystems.

## Author contributions

CSY funded the project, led the writing of the manuscript, contributed tidal data, contributed to interpretation, and assisted in field work. JBG devised the concept, its analysis and contributed to its interpretation and writing. YJC assisted in fieldwork, organized and analysed the data. YY analysed and provided the wind field data and contributed to the interpretation. DC conducted the fieldwork.

## Acknowledgements

The authors thank the supporting staff at the Centre for Marine and Coastal Studies, Universiti Sains Malaysia, for help on field work and Dr. Foong Swee Yeok (School of Biological Sciences, Universiti Sains Malaysia) and Dr. Aldrie Amir (Universiti Kebangsaan Malaysia) for their suggestions and advice during sampling and writing.

## Data reference

Chee, Su Yin; Gallagher, John Barry; Yusup, Yusri; Yee, Jean Chai; Carey, Danielle (2019), “Anthropogenic marine debris found in peri-urban and urban mangroves on Penang Island, Malaysia”, Mendeley Data, v1 http://dx.doi.org/10.17632/7tc9j8nynb.1

## Notes

http://dx.doi.org/10.17632/7tc9j8nynb.1

